# Identification of putative druggable pockets in PRL3, a significant oncology target, using *in silico* analysis

**DOI:** 10.1101/2023.07.31.551065

**Authors:** Grace M. Bennett, Julia Starczewski, Mark Vincent C. dela Cerna

## Abstract

Due to their active roles in regulating phosphorylations, protein tyrosine phosphatases (PTP) have emerged as attractive targets in diseases characterized by aberrant phosphorylations such as cancers. The activity of the phosphatase of regenerating liver 3, PRL3, has been linked to several oncogenic and metastatic pathways, particularly in breast, ovarian, colorectal, and blood cancers. Development of small molecules that directly target PRL3, however, has been challenging. This is partly due to the lack of structural information on how PRL3 interacts with its inhibitors. Here, computational methods are used to bridge this gap by evaluating the druggability of PRL3. In particular, web-based pocket prediction tools, DoGSite3 and FTMap, were used to identify binding pockets using structures of PRL3 currently available in the Protein Data Bank. Druggability assessment by molecular dynamics simulations with probes was also performed to validate these results and to predict the strength of binding in the identified pockets. While several druggable pockets were identified, those in the closed conformation show more promise given their volume and depth. These two pockets flank the active site loops and roughly correspond to pockets predicted by molecular docking in previous papers. Notably, druggability simulations predict the possibility of low nanomolar affinity inhibitors in these sites implying the potential to identify highly potent small molecule inhibitors for PRL3. Putative pockets identified here can be leveraged for high-throughput virtual screening to further accelerate the drug discovery against PRL3 and development of PRL3-directed therapeutics.

## Introduction

Aberrant cellular phosphorylation is a hallmark of several diseases including inflammation and cancers (*1*–*3*). Protein phosphorylation is a post-translational modification that can act as a switch to regulate biochemical pathways and is regulated by the concerted action of two classes of enzymes: kinases and phosphatases. As such, kinases and phosphatases present as significant potential clinical molecular targets. To date, several kinase inhibitors have been approved for various indications and kinases have been regarded as one of the most important drug targets of the 21^st^ century (*4*–*8*). Meanwhile, phosphatases are recently gaining traction as therapeutic drug targets, particularly with emerging roles in cancers (*9*–*12*). Development of phosphatase inhibitors offer a novel approach to treatment of diseases involving dysregulated protein phosphorylation.

Among protein phosphatases, the phosphatase of regenerating liver (PRL), also known as the protein tyrosine phosphatase 4A (PTP4A), family is of significant interest in drug discovery as PRL expression has been correlated with oncogenicity (*13*–*16*). PRL3 or PTP4A3 is the most well-studied PRL and is highly expressed in several cancer types. Its expression has been correlated with poor patient prognosis in various cancers as well as with increased proliferation and metastatic potential in cellular models (*17*–*26*). While a lot remains unknown about its physiological functions and endogenous substrates, PRL3 has been shown to be involved in regulation of apoptosis, cellular metabolism, DNA integrity, epithelial-to-mesenchymal transition (EMT), and angiogenesis (*27*). As such, PRL3 is emerging as one of the most prominent phosphatase drug targets in recent years.

Given its roles in cancers, PRL3 has been the target of several drug discovery programs leading to the identification of several potential inhibitors (*11, 13, 14, 28*–*35*). Pentamidine isethionate, an FDA approved anti-protozoa drug, is among the first molecules to be identified to inhibit PRLs *in vitro* (*13*). High-throughput screening efforts identified several potential PRL3 inhibitors including several rhodanine derivatives, and one of the most potent inhibitors at the time, thienopyridone (*29*–*32, 34, 36*). Further elaboration of the thienopyridone scaffold eventually led to the development of JMS-053, currently the most potent experimental inhibitor of PRL3 (*33, 37, 38*). Recently, the JMS-053 scaffold was linked to an adamantly moiety to generate a bifunctional ER stress inducer/PRL3 inhibitor (*35*). Despite the success of these campaigns to identify candidate molecules, however, no small molecule inhibitor targeting PRL3 has advanced to clinical studies. The rhodanine scaffold has been considered a promiscuous binder and likely is associated with several off-target effects (*37, 39*). Meanwhile, the thienopyridone scaffold exhibits redox activity which potentially inhibits enzymes susceptible to oxidation, such as PRL3 (*37, 40*). Thus, while there a few candidate molecules, the search for inhibitors for PRL3 with potential to be developed as cancer therapeutics remains open.

To date, a few structures of PRL3 have been experimentally determined, capturing its open and closed conformations (*41*–*44*). The closed conformation was determined in the presence of vanadate, a general protein tyrosine phosphatase inhibitor, bound to the active site (*44, 45*). A crystal structure of PRL3 has also been determined where the CBS-pair domain of the magnesium transporter CNNM3 is bound to the active site (*41*). In this interaction, PRL3 acts as a pseudo-phosphatase, revealing a unique cellular function for the PRL3 family. While these structures have allowed for characterization of function of PRL3, a challenge remains in that none of these structures capture how PRL3 binds to small molecules, which is critical for structure-based drug design and virtual screening campaigns.

In this present work, available structural information is leveraged to identify and characterize putative druggable pockets within PRL3. The objective is to inform high throughput *in silico* screening efforts by focusing on pockets amenable for inhibitor binding. Pocket identification tools were used, along with molecular dynamics simulations using probes, to identify druggable pockets towards the development of highly specific PRL3 inhibitors.

## Methods

### Protein Structures

All analyses were performed on three structures of PRL3 taken from the Protein Data Bank: 2MBC, 1V3A, and 5TSR. These structures represent the open (1V3A) and closed (2MBC, 5TSR) conformations of PRL3 (*41, 44*). One of the closed conformations (5TSR) is the structure of PRL3 bound to the CNNM3 magnesium transporter Bateman domain determined by X-ray crystallography to a resolution of 3.19 Å. Chain A, the PRL3 structure, was extracted from this. A closed conformation in the presence of vanadate (2MBC) was determined by solution NMR. Only the first model of the twenty in the NMR bundle was used. The open conformation (1V3A) was also determined by solution NMR and only a single conformer was deposited (*42*). All these structures were used without further modification.

### Druggability simulations and analysis

Druggability molecular dynamics simulations were set-up using the druggability suite VMD plug-in (DruGUI) and were performed using the NAMD software and CHARMM forefield (*46, 47*). PRL3 structures were placed in a box with 8 Å padding, containing explicit TIP3P water and enough chloride ions to neutralize the system. The probe composition for all runs was set at 80% isopropanol and 20% acetamide.

Simulations were first minimized for 2000 steps prior to a series of equilibration steps. First, the system was heated from 100 K to 300 K over 40 ps and ran at 300 K for 80 ps. Then, the system was further heated to 600 K over 60 ps, ran at 600 K for 600 ps, and cooled back to 300 K over 18 ps. The Cα atoms were restrained by a harmonic potential with a force constant of 1 kcal/mol/A^2^ during these equilibration steps. A final 600 ps equilibration without constraint was done at 300 K. Two independent 40-ns production runs were carried out for each of the systems.

Analysis of probe binding hotspots was done using the built-in analysis tool in the DruGUI plug-in. The default parameters were used for the analyses of all simulations. The two simulations for each system were analyzed individually. Results were visualized using VMD, PyMol, or ChimeraX (*48*–*50*).

### Detection of possible binding pockets

Two web/server-based tools were used to detect and identify potentially druggable pockets. The tools were chosen as they employ distinct algorithms in identifying protein pockets and are both free to use. DoGSite3 is an automated grid-based pocket detection tool included in the Protein*Plus* server (Center of Bioinformatics, University of Hamburg). It uses Difference of Gaussian (DoG) filter for pocket prediction and considers volume, hydrophobicity, enclosure, and depth (*51*–*53*). Meanwhile, FTmap relies on the identification of binding hotspots of 16 small chemical probes. Docked probes go through energy-based clustering followed by consensus clustering to identify binding hotspots (*54*). The prepared structures were directly uploaded onto these web tools and results were visualized with either PyMol or ChimeraX (*48, 49*).

## Results and Discussion

Protein tyrosine phosphatases are now emerging as significant therapeutic targets, particularly in cancers (*3, 55*). Among the PRL sub-family, PRL3 is the most well-studied and most targeted by on-going drug discovery efforts. While there have been several promising inhibitors identified and structures of PLR3 experimentally determined, there is currently no information on how PRL3 interacts with any of these small molecules. That is, there is currently no known structure of PRL3 in complex with any inhibitor. Information on the interaction of a protein target and inhibitors is critical for the rational improvement of these inhibitors and the design of new ones (*56*). In the absence of these structures, any information on putative binding pockets is crucial to *in silico* drug screening efforts (*51, 54, 56, 57*). This study aims to evaluate the druggability of PRL3 and to identify and characterize putative druggable pockets which will further inform drug discovery and development against this important target. To maximize the likelihood of identifying a druggable pocket, multiple structures (open, closed, and pseudo-substrate-bound, Supplementary Figure 1) are used along with multiple computational tools.

Two web/server-based binding pocket detection tools with unique detection algorithms were used, along with molecular dynamics druggability simulations.

DoGSite3 is an improvement on DoGSiteScorer that better handles binding site boundary using a depth-first search (*51*). It successfully identified several putative pockets in all three conformations of PRL3 used (Supplementary Tables 1 and 2, Figure 1). Five putative binding pockets were identified in the open conformation ranging from 46 to 127 Å^3^ in volume (Figure 1A, C). The 127 Å^3^ pocket has a depth of 7.6 Å and is the largest in this conformation. The vanadate-bound, closed conformation has larger pockets with two prominent one having volumes of 221 Å^3^ and 181 Å^3^ (Figure 1B, C). These are adjacent to the active site loops, the WPD and P loops, respectively. These pockets are also the deepest ones detected at 13.6 Å for the WPD-adjacent and 10.3 Å and P-adjacent pockets (Figure 1D, E). These pockets involve the active site residues D72, C104, and R110 and are highly hydrophobic (64% and 84% hydrophobic residues, respectively, Figure 1D, E, Supplementary Table 1). The pseudo-substrate-bound structure, interestingly, did not show comparable binding pockets (Supplementary Table 1). Interestingly, using the older version of DoGSite3, two adjacent pockets that cover the entire active site are identified, in addition to several others, with a combined volume of 1,030 Å^3^ (Supplementary Table 2). This is not surprising considering that PRL3 has an active site that is more shallow than typical PTP active sites and DoGSite3 factors in the depth of the pocket (*42, 43*).

**Figure 1.**
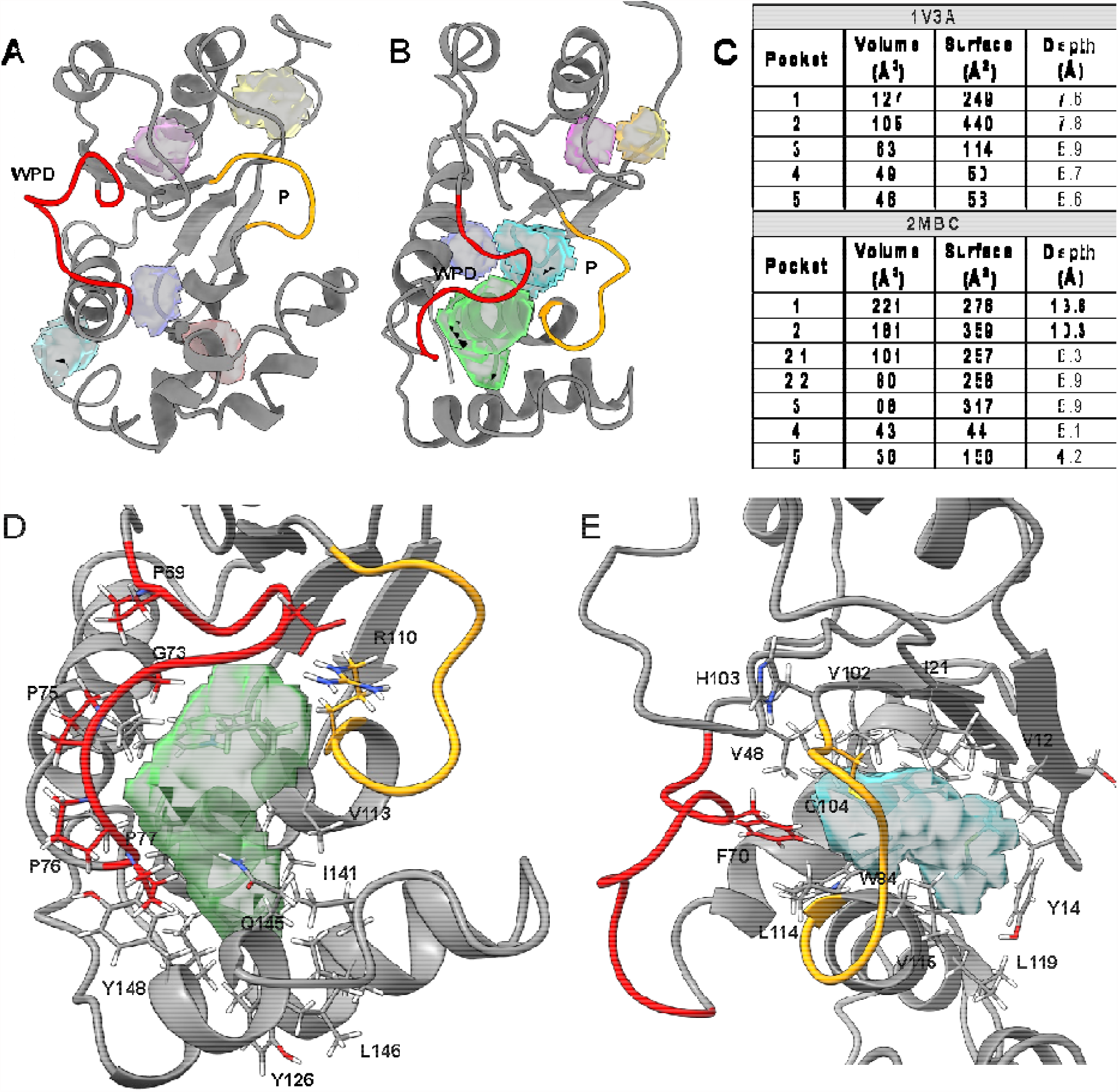
Pockets identified by DoGSite3. Several small pockets were identified in the open (1V3A, A, C) and closed conformations (2MBC, B, C) of PRL3. Pocket parameters, including volume, surface, and depth are summarized (C, Supplementary Table 1). Two major pockets are identified in the closed conformation (Pockets 1 and 2 in 2MBC) cradled near the active site loops (red and orange).

Another approach to identifying putative binding pockets in a protein is the computational analogue to fragment screening (*58, 59*). One such implementation is FTmap, which performs rigid docking of 16 small molecule probes to identify clusters or hotspots which correspond to possible drug binding pockets (*54*). Using FTmap, several potential binding pockets were identified in all three conformations studied, some of which overlap with pockets identified by DoGSite3 (Figure 2). While the active site in the open conformation does not meet the depth criteria of DoGSite3, pockets identified by DoGSiteScorer within the active site overlap with FTmap hotspots (Supplementary Figure 2). Fragment binding predicted by FTmap may imply that, while shallow, it is still possible to design molecules that bind to this pocket. Interestingly, a small pocket near the N-terminus also overlaps with an FTmap hotspot (Figure 2A). Meanwhile, for the vanadate-bound conformation, the biggest FTmap clusters of probes overlap with the biggest pockets identified by DoGSite3, adjacent to the WPD and P loops (Figure 2B). Similarly, several hotspots were identified for the pseudo-substrate-bound conformation, including some that overlap with the smaller DoGSite3 pockets (Figure 2C). In addition to identifying hotspots from consensus clusters based on docked probes, FTmap also quantifies the non-bonded and hydrogen bonding interaction between residues and the probes (Figure 3, Supplementary Figure 3). For the vanadate-bound closed conformation, there is significant involvement from the active site loops (WPD and P) in binding the hotspots (Figure 3A, B). The major interacting residues (15% of the top residue count or higher) map to residues that are identified in DoGSite3 as well (Figure 3C, D). Moreover, there is significant non-bonded interactions identified near the N-terminus (residues 10-20, Figure 3B) which is potentially another pocket, albeit significantly smaller. This area was also identified by DogSite3 (Figure 1B). Active site involvement is not as prominent in the open conformation, although there is potential for significant interactions adjacent to the P loop (residues 110-120, Supplementary Figure A, B). This is again reflective of the shallow binding pocket of PRL3; though interestingly, several hotspots are detected in that shallow pocket (Figure 2A).

**Figure 2.**
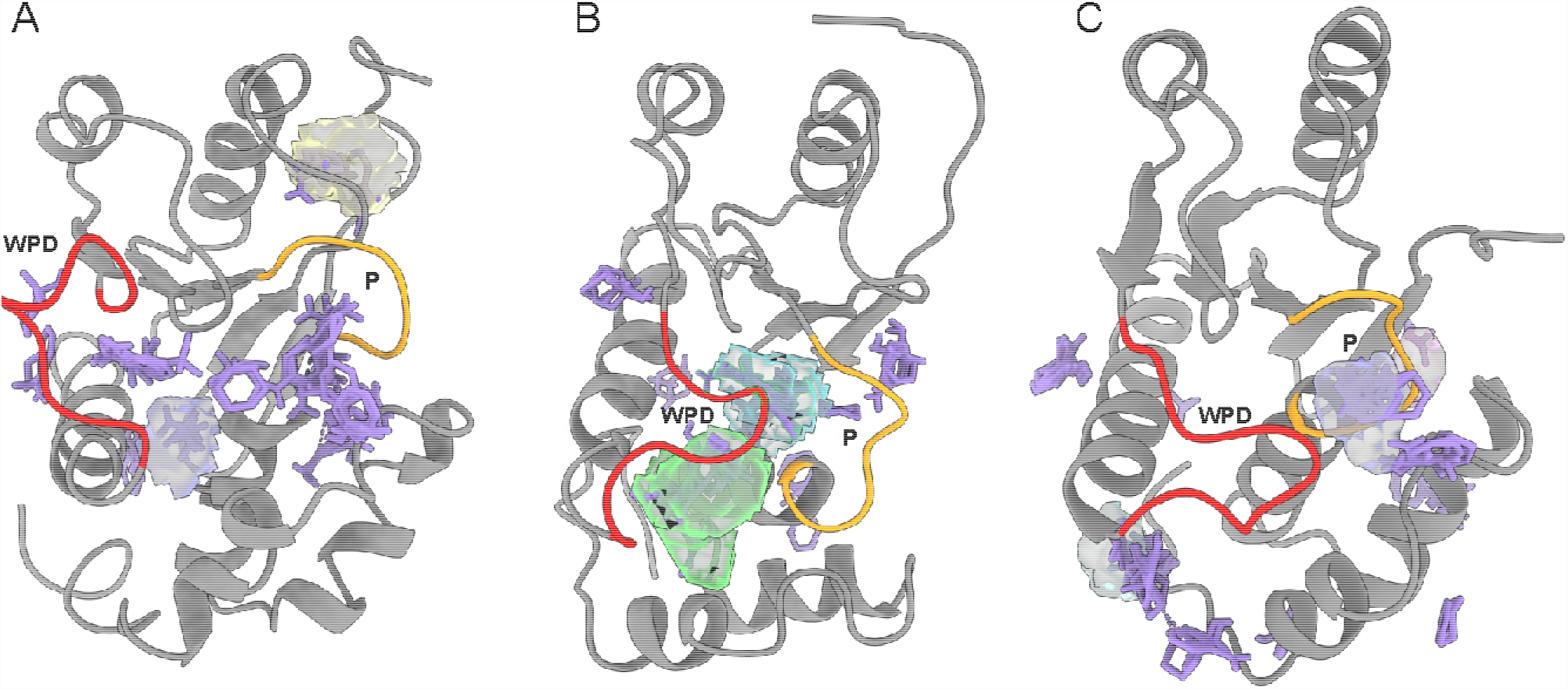
Fragment hotspots determined by FTmap. Several hotspots for the various probes (all colored purple to emphasize positions regardless of probe identity) were identified in the open (1V3A, A), vanadate-bound (2MBC, B), and pseudo-substrate bound (5TSR, C) conformations. Where there is overlap, the DoGSite3-predicted pockets are also shown.

**Figure 3.**
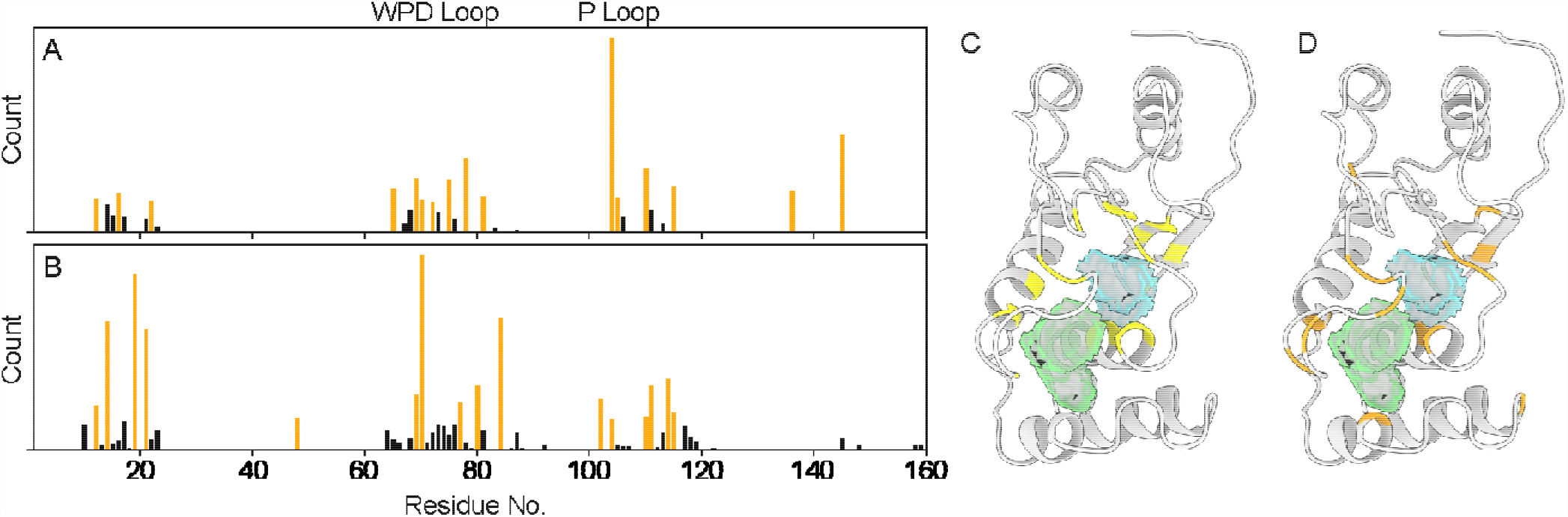
Residue level interaction data from FTmap. Raw counts of hydrogen bonding (A) and non-bonded (B) interactions in the closed conformation (2MBC). Active site loops are highlighted (WPD, yellow and P, orange) along with significant interaction counts, defined as 15% of the highest counts or more (orange bars). These top hydrogen bonding (C) and non-bonding (D) interactions are mapped onto the structure along with the primary pockets identified in DoGSite3 (yellow and blue surfaces).

Overall, these two approaches identified several pockets of interest within PRL3. It is noteworthy that based on these two techniques, the vanadate-bound closed conformation reveals the most promising binding pockets (Figure 1D, E, 2B, Supplementary Table 1). These pockets have sufficient volume and depth to be identified by DoGSite3 and several clusters/hotspots and interaction data determined by FTmap support this. There are also some sites in the open conformation identified by both methods, although they are significantly smaller (Figure 2A). The difference in pocket identification algorithm also allows FTmap to identify potential hotspots within the active site of the open conformation that is missed by DoGSite3 (Figure 2A, Supplemental Figure 2). While these pockets are capable of hydrogen bonding, majority of the residues are hydrophobic (Supplementary Table 2).

To further analyze these pockets, a molecular dynamics druggability simulation was performed using the DruGUI VMD plugin (*47, 50*). This offers yet another unique approach to validate the results so far. Like FTmap, druggability simulations makes use of small molecule probes to identify potential druggable pockets (*54*). A molecular dynamics simulation in explicit water in the presence of molecular probes is used to identify hotspots and estimate maximal predicted binding affinity at a site, as opposed to the rigid docking method (*47*). Several hotspots were identified by the druggability simulations (Supplementary Table 3), particularly in the open and vanadate-bound closed conformation. The top hotspots, based on predicted lowest binding energy (or highest binding affinity) for the open and vanadate-bound conformations capture some of the previously identified pockets (Table 2). Several hotspots have highly desirable predicted maximum binding affinities from low nanomolar to <100 μM, indicating that these hotspots might be promising druggable pockets. The top predicted binding pocket in the open conformation, for instance, is the shallow active site pocket (Figure 4A) with a theoretical strongest binding affinity at the nanomolar range. A similarly potentially high affinity binding pocket is detected in the closed conformation, adjacent to the P loop (Figure 4B). This site roughly corresponds with the P loop-adjacent site identified by DoGSite3 and FTmap (Figure 2B, 3).

**Table 1.** Results from DoGSite3.

| **2MBC** |  |  |  |
| --- | --- | --- | --- |
| **Pocket** | **Volume**<br>**(Å<sup>3</sup>)** | **Surface**<br>**(Å<sup>2</sup>)** | **Depth**<br>**(Å)** |
| P1 | 221 | 276 | 13.6 |
| P2 | 181 | 359 | 10.3 |
| P2_1 | 101 | 257 | 8.3 |
| P2_2 | 80 | 258 | 6.9 |
| P3 | 68 | 317 | 5.9 |
| P4 | 43 | 44 | 5.1 |
| P5 | 36 | 156 | 4.2 |
| **1V3A** |  |  |  |
| **Pocket** | **Volume**<br>**(Å<sup>3</sup>)** | **Surface**<br>**(Å<sup>2</sup>)** | **Depth**<br>**(Å)** |
| P1 | 127 | 249 | 7.6 |
| P2 | 105 | 440 | 7.8 |
| P3 | 63 | 114 | 5.9 |
| P4 | 49 | 50 | 5.7 |
| P5 | 46 | 56 | 5.6 |
| **5TSR** |  |  |  |
| **Pocket** | **Volume**<br>**(Å<sup>3</sup>)** | **Surface**<br>**(Å<sup>2</sup>)** | **Depth**<br>**(Å)** |
| P1 | 81 | 275 | 6.9 |
| P2 | 60 | 204 | 5.6 |
| P3 | 47 | 146 | 5.8 |
| P4 | 26 | 80 | 4.4 |

**Table 2.**
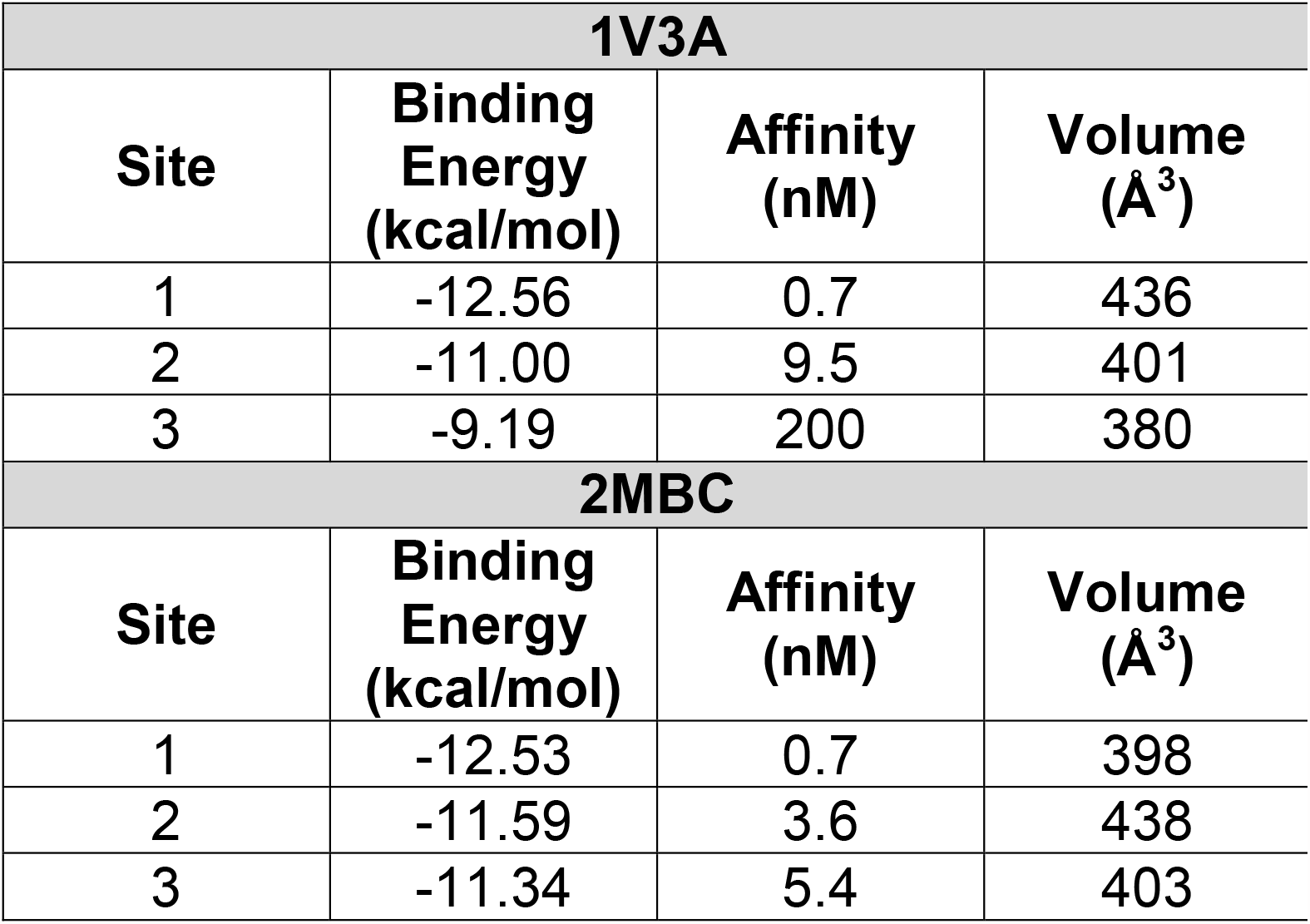
Hotspots predicted by druggability simulations.

| **1V3A** |  |  |  |
| --- | --- | --- | --- |
| **Site** | **Binding Energy (kcal/mol)** | **Affinity (nM)** | **Volume (Å<sup>3</sup>)** |
| 1 | -12.56 | 0.7 | 436 |
| 2 | -11.00 | 9.5 | 401 |
| 3 | -9.19 | 200 | 380 |
| **2MBC** |  |  |  |
| **Site** | **Binding Energy (kcal/mol)** | **Affinity (nM)** | **Volume (Å<sup>3</sup>)** |
| 1 | -12.53 | 0.7 | 398 |
| 2 | -11.59 | 3.6 | 438 |
| 3 | -11.34 | 5.4 | 403 |

**Figure 4.**
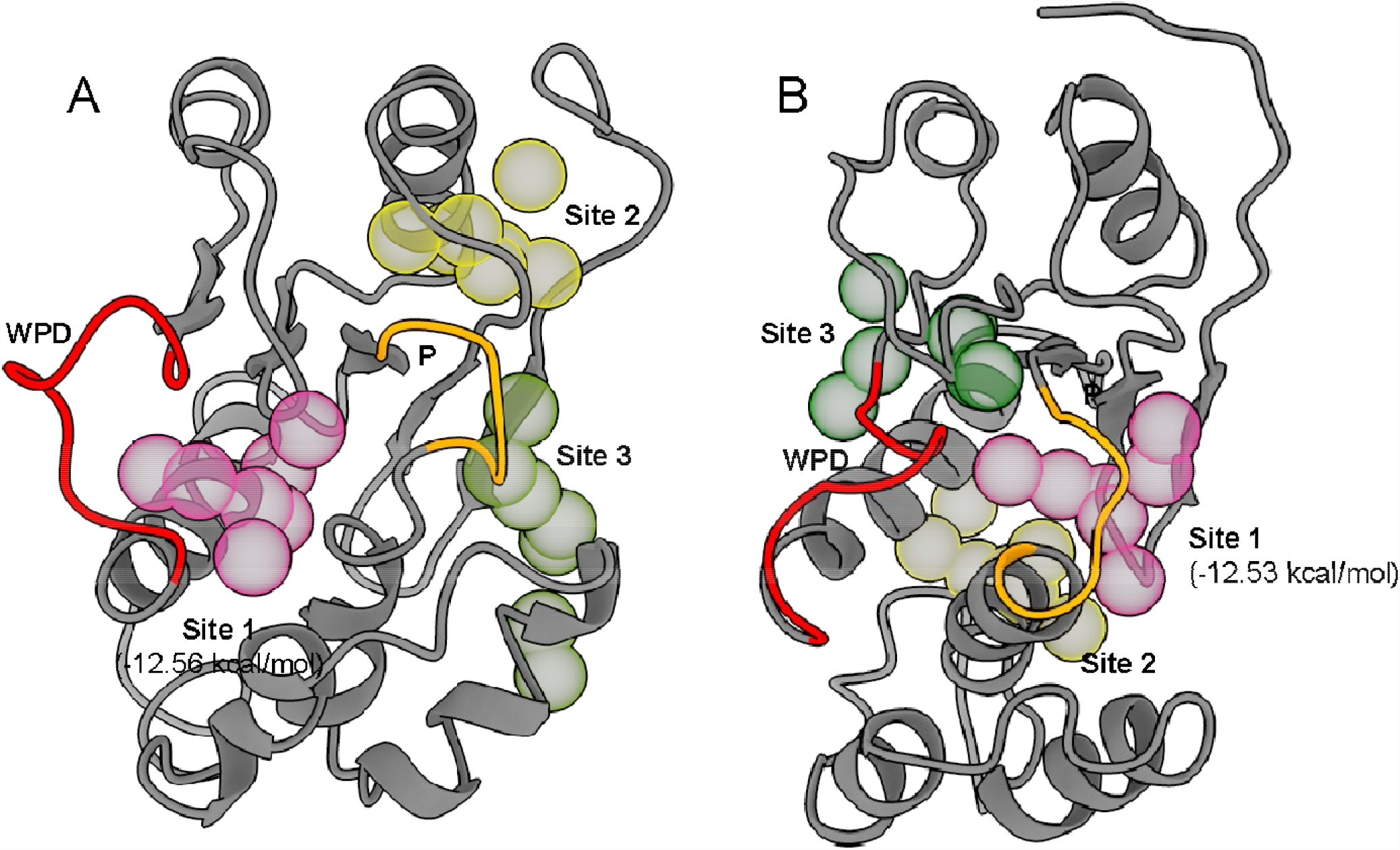
Druggability simulations identify probe hotspots. Top 3 hotspots from DruGUI druggability simulations for the open (1V3A, A) and vanadate-bound closed (2MBC, B) conformation of PRL3.

Overall, this study has identified several potentially druggable binding pockets, including the active site and most notably a binding pocket in the closed conformation that is adjacent to the active site P loop. Residues that line these pockets have also been identified for use in high-throughput flexible side chain docking simulations (*62*–*64*). Druggability simulations predict high affinity binders in several pockets. While this is a theoretical maximum, this provides hope for future drug discovery programs to identify potent PRL3 inhibitors. The pockets identified here also support and validate previous predictions. For instance, a previous virtual screening attempt identified inhibitors that bind the shallow active site (*32*). The thienopyridone scaffold was also docked in the closed conformation near the WPD loop, near one of the sites identified in the present study (*38*). Blind docking of FDA-approved drugs similarly identified roughly the same pockets in both the open and closed conformations (*60*). In the absence of experimental structures of PRL3 in complex with inhibitors, these computational studies provide invaluable information that will guide drug design efforts against this important therapeutic target.

## Conclusion

PRL3 is a protein tyrosine phosphatase (PTP) that has emerged as a significant oncology drug target. While PTPs have historically been labelled ‘undruggable,’ this family of proteins is slowly shedding this identity (*9*–*12, 61*). This study contributes to further advancing PRL3-targeted drug discovery by predicting potentially druggable pockets within the open and closed conformations. Three unique methodologies identified some consensus sites, as well as unique sites, that could be the focus of virtual drug screening, using the vast chemical space accessible. Knowledge on the residues that might be involved in drug binding can now be used in conjunction with docking with flexible sidechains (*62*–*64*). Furthermore, this study contributes to further supporting the druggability of PRL3 – that perhaps more high affinity inhibitors are just soon to be discovered.

## Acknowledgements

GB is funded by the American Society for Biochemistry and Molecular Biology Undergraduate Research Award. MVCDC acknowledges support from the Department of Biochemistry, Chemistry, and Physics and the College of Science and Mathematics at Georgia Southern University in the form of a start-up package.

