## Supplementary Tables and Figures for "Identification of putative druggable pockets in PRL3, a significant oncology target, using *in silico* analysis"

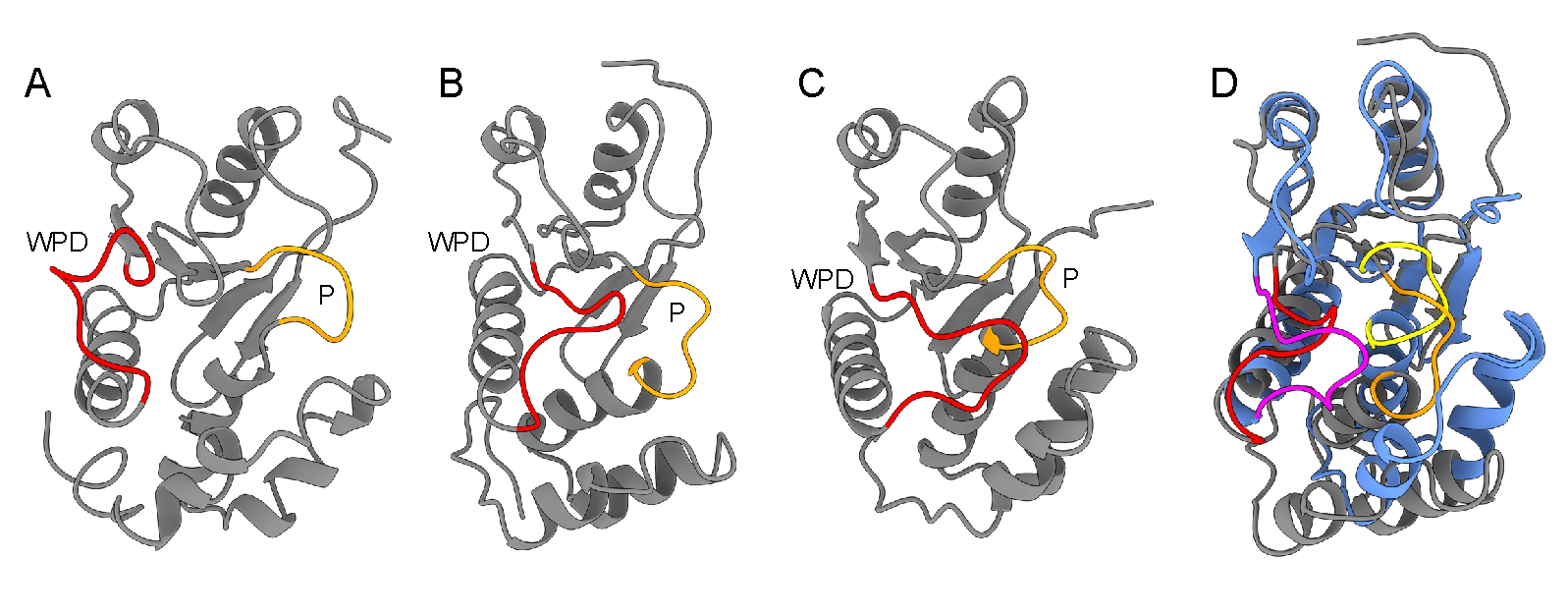

**Supplemental Figure 1. Structures of PRL3.** The open (1V3A, A) and closed (2MBC, B and 5TSR, C) conformations of PRL3 have been experimentally determined and exhibit the opening and closing of the active site, WPD and P, loops. The structure of PRL3 bound to a pseudo-substrate, the CBS-pair domain of CNNM3, (C) is distinct from the vanadate-bound closed conformation (B), particularly in the positioning of the active site loops (WDP: magenta vs red, P: yellow vs orange, D).

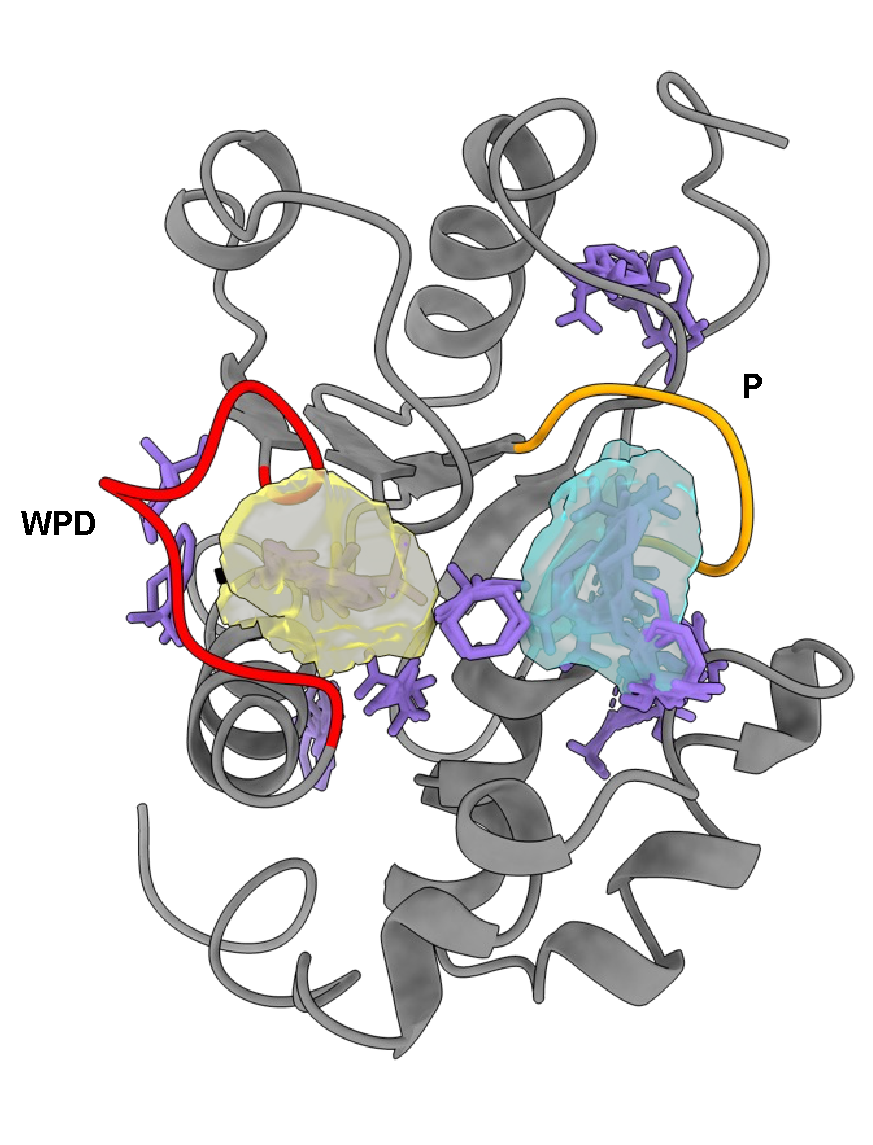

**Supplemental Figure 2. Shallow binding pocket in the open conformation of PRL3.** Several FTmap hotspots within the active site of PRL3 fall within pocket P1_1 (yellow) and P3 (cyan) identified by DoGSiteScorer (Supplementary Table 2).

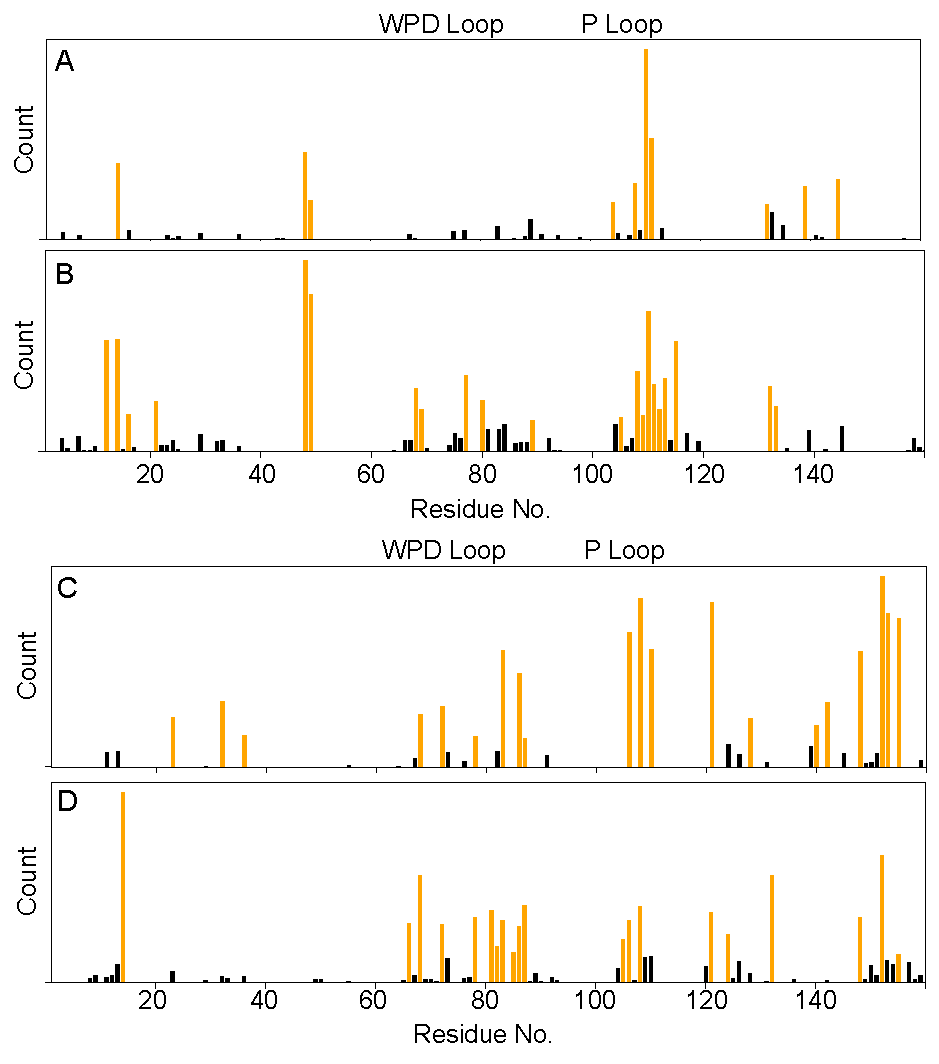

**Supplementary Figure 3. Residue level interaction data from FTmap.** Hydrogen bonding (A) and non-bonded (B) interaction counts for the open conformation (1V3A). Hydrogen bonding (C) and non-bonded (D) interaction counts for the pseudo-substrate-bound conformation (5TSR).

**Supplementary Table 1. Binding Pocket Prediction by DoGSite3**

| **1V3A** | | | | |
| --- | --- | --- | --- | --- |
| **Pocket** | **Volume (Å^3^)** | **Surface (Å^2^)** | **Depth (Å)** | **Residues** |
| 1 | 127 | 249 | 7.6 | A2, M4, N5, P7, A8, P9, T22, H23, N24, P25, T29, T32, F33, D36 |
| 2 | 105 | 440 | 7.8 | E82, L85, S86, K89, Y152, Q156, R157, L158, K161, D162 |
| 3 | 63 | 114 | 5.9 | V48, W68, P77, V80, V81, W84, V113, L114, L117 |
| 4 | 49 | 50 | 5.7 | T26, L30, F33, V45, V46, R47, Y53, V65, V66, A101, V102, H103 |
| 5 | 46 | 56 | 5.6 | P112, V113, A116, L117, I120, Y126, A129, I130, I133, Q145, L146, Y148 |
| **2MBC** | | | | |
| **Pocket** | **Volume (Å^3^)** | **Surface (Å^2^)** | **Depth (Å)** | **Residues** |
| 1 | 221 | 276 | 13.6 | P69, F70, D72, G73, A74, P75, P76, P77, G78, V80, V81, W84, R110, V113, L114, A116, L117, I120, Y126, I130, I141, Q145, L146, Y148, L149 |
| 2 | 181 | 359 | 10.3 | V12, S13, Y14, M17, F19, I21, V48, F70, W84, L87, V88, V102, H103, C104, A111, L114, V115, A118, L119 |
| 2.1 | 101 | 257 | 8.3 | V12, F19, I21, V48, F70, W84, L87, V102, H103, C104, A111, L114, V115 |
| 2.2 | 80 | 258 | 6.9 | V12, S13, Y14, M17, F19, W84, V88, V115, A118, L119 |
| 3 | 68 | 317 | 5.9 | K15, H16, M17, F92, S122, L158, R159 |
| 4 | 43 | 44 | 5.1 | P9, L20, I21, T22, F33, D36, L37, Y40, V46, A101 |
| 5 | 36 | 156 | 4.2 | P7, A8, P9, T22, H23, N24, N27, D36 |
| **5TSR** | | | | |
| **Pocket** | **Volume (Å^3^)** | **Surface (Å^2^)** | **Depth (Å)** | **Residues** |
| 1 | 81 | 275 | 6.9 | M17, V88, K89, F92, C93, L158, R159 |
| 2 | 60 | 204 | 5.6 | V81, E82, L85, E121, Y152, R153, P154, K155, R157 |
| 3 | 47 | 146 | 5.8 | D72, G73, A104, V105, A106, L108, G109, R110, N142, Q145 |
| 4 | 26 | 80 | 4.4 | V10, V12, I21, G107, P112, I133, K136, R137, A140 |

**Supplementary Table 2. Binding Pocket Prediction by DoGSiteScorer**

| **2MBC** | | | | |
| --- | --- | --- | --- | --- |
| **Pocket** | **Volume (Å^3^)** | **Surface (Å^2^)** | **Drug Score** | **Simple Score** |
| P0 | 1022 | 1170 | 0.82 | 0.69 |
| P0_0 | 562 | 739 | 0.72 | 0.60 |
| P0_1 | 460 | 530 | 0.71 | 0.47 |
| P1 | 259 | 569 | 0.66 | 0.16 |
| P1_0 | 155 | 416 | 0.28 | 0.14 |
| P1_1 | 104 | 220 | 0.28 | 0.02 |
| P2 | 248 | 329 | 0.62 | 0.06 |
| P3 | 133 | 397 | 0.19 | 0.00 |
| **1V3A** | | | | |
| **Pocket** | **Volume (Å^3^)** | **Surface (Å^2^)** | **Drug Score** | **Simple Score** |
| P0 | 1233 | 1720 | 0.82 | 0.62 |
| P0_0 | 332 | 545 | 0.4 | 0.35 |
| P0_1 | 282 | 502 | 0.36 | 0.26 |
| P0_2 | 253 | 365 | 0.51 | 0.26 |
| P0_3 | 129 | 216 | 0.33 | 0.05 |
| P0_4 | 101 | 193 | 0.47 | 0.00 |
| P0_5 | 81 | 265 | 0.08 | 0.00 |
| P0_6 | 55 | 202 | 0.13 | 0.00 |
| P1 | 620 | 922 | 0.86 | 0.48 |
| P1_0 | 323 | 496 | 0.72 | 0.36 |
| P1_1 | 175 | 332 | 0.39 | 0.13 |
| P1_2 | 123 | 327 | 0.25 | 0.07 |
| P2 | 410 | 671 | 0.78 | 0.26 |
| P2_0 | 204 | 434 | 0.2 | 0.15 |
| P2_1 | 146 | 274 | 0.45 | 0.12 |
| P2_2 | 60 | 117 | 0.67 | 0.00 |
| P3 | 164 | 299 | 0.36 | 0.00 |
| **5TSR** | | | | |
| **Pocket** | **Volume (Å^3^)** | **Surface (Å^2^)** | **Drug Score** | **Simple Score** |
| P0 | 351 | 708 | 0.65 | 0.25 |
| P1 | 183 | 319 | 0.54 | 0.00 |
| P1_0 | 72 | 171 | 0.14 | 0.00 |
| P1_1 | 61 | 105 | 0.36 | 0.00 |
| P1_2 | 50 | 225 | 0.17 | 0.00 |
| P2 | 151 | 372 | 0.35 | 0.00 |
| P3 | 143 | 205 | 0.32 | 0.00 |
| P4 | 119 | 277 | 0.46 | 0.00 |
| P5 | 115 | 254 | 0.25 | 0.00 |

**Supplementary Table 3. DruGUI binding energy and predicted affinities**

| **1V3A** | | | | |
| --- | --- | --- | --- | --- |
| **Site** | **Solution** | **Binding Energy (kcal/mol)** | **Affinity (nM)** | **Volume (Å^3^)** |
| 1 | Sim1-1 | -11.24 | 6.4 | 432 |
|  | Sim1-2 | -11.22 | 6.6 | 451 |
|  | Sim1-3 | -11.10 | 8.1 | 445 |
|  | Sim2-1 | -12.56 | 0.7 | 436 |
|  | Sim2-2 | -12.50 | 0.8 | 430 |
|  | Sim2-3 | -12.43 | 0.9 | 408 |
| 2 | Sim1-1 | -10.52 | 21.6 | 420 |
|  | Sim1-2 | -10.50 | 22 | 431 |
|  | Sim1-3 | -10.49 | 22.7 | 438 |
|  | Sim2-1 | -11.00 | 9.5 | 401 |
|  | Sim2-2 | -10.90 | 11.4 | 425 |
|  | Sim2-3 | -10.85 | 12.2 | 411 |
| 3 | Sim1-1 | -9.19 | 200 | 380 |
|  | Sim1-2 | -9.10 | 231 | 400 |
|  | Sim1-3 | -9.09 | 237 | 417 |
|  | Sim2-1 | -8.17 | 1100 | 388 |
| 4 | Sim2-1 | -9.42 | 136 | 409 |
|  | Sim2-2 | -9.28 | 171 | 430 |
|  | Sim2-3 | -9.23 | 186 | 413 |
| **2MBC** | | | | |
| **Site** | **Solution** | **Binding Energy (kcal/mol)** | **Affinity (nM)** | **Volume (Å^3^)** |
| 1 | Sim1-1 | -12.53 | 0.7 | 398 |
|  | Sim1-2 | -12.28 | 1.1 | 398 |
|  | Sim1-3 | -12.23 | 1.2 | 417 |
| 2 | Sim2-1 | -11.59 | 3.6 | 438 |
|  | Sim2-2 | -11.54 | 3.9 | 416 |
|  | Sim2-3 | -11.54 | 3.9 | 447 |
| 3 | Sim2-1 | -11.34 | 5.4 | 403 |
|  | Sim2-2 | -11.27 | 6.1 | 425 |
|  | Sim2-3 | -11.25 | 6.3 | 368 |
| **5TSR** | | | | |
| **Site** | **Solution** | **Binding Energy (kcal/mol)** | **Affinity (nM)** | **Volume (Å^3^)** |
| 1 | Sim1-1 | -10.05 | 47.3 | 426 |
|  | Sim1-2 | -9.47 | 125 | 442 |
| 2 | Sim1-1 | -9.69 | 85.9 | 356 |
| 3 | Sim1-1 | -9.62 | 97.5 | 350 |
